# Plant trichomes and a single gene *GLABRA1* contribute to insect community composition on field-grown *Arabidopsis thaliana*

**DOI:** 10.1101/320903

**Authors:** Yasuhiro Sato, Rie Shimizu-Inatsugi, Misako Yamazaki, Kentaro K. Shimizu, Atsushi J. Nagano

## Abstract

**Background:** Genetic variation in plants alters insect abundance and community structure in the field; however, little is known about the importance of a single gene among diverse plant genotypes. In this context, *Arabidopsis* trichomes provide an excellent system to discern the roles of natural variation and a key gene, *GLABRA1*, in shaping insect communities. In this study, we transplanted two independent glabrous mutants (*gl1-1* and *gl1-2*) and 17 natural accessions of *Arabidopsis thaliana* to two localities in Switzerland and Japan.

**Results:** Fifteen insect species inhabited plant accessions, with 10–30% broad-sense heritability of community indices being detected, such as species richness and diversity. The total abundance of leaf-chewing herbivores was negatively correlated with trichome density at both the field sites, while glucosinolates had variable effects on leaf chewers between the two sites. Interestingly, there was a parallel tendency for the abundance of leaf chewers to be higher on *gl1-1* and *gl1-2* than for their different parental accessions, L*er*-1 and Col-0, respectively. Furthermore, the loss of function in the *GLABRA1* gene significantly decreased the resistance of plants to the two predominant chewers, flea beetles and turnip sawflies.

**Conclusions:** Overall, our results indicate that insect community composition on *A. thaliana* is heritable across two distant field sites, with *GLABRA1* playing a key role in altering the abundance of leaf-chewing herbivores. Given that such a trichome variation is widely observed in Brassicaceae plants, the present study exemplifies the community-wide impact of a single plant gene on crucifer-feeding insects in the field.

## Background

Plants develop various resistance traits, such as spines and toxins, to deter herbivory [1]. A growing number of studies on community genetics has revealed that genetic variation in plant resistance traits exerts cascading effects on insect abundance and community composition [2, 3, 4, 5]. These insect indices projected on individual plants, called extended phenotype [5], exhibit heritable variation among plant genotypes [6, 7, 8]. Some researchers have reported the association of particular genetic polymorphisms with leaf damage [9, 10], insect abundance [3, 11], and community composition [3] in the field. In comparison, other studies have focused on how single genes affect the insect community using transformed plants [12, 13]. These lines of evidence from diverse plant species suggest that both quantitative genetic variation and single genes contribute to the community genetics of plant-insect interaction.

*Arabidopsis thaliana* L. (Brassicaceae) is well-studied as a model system of Brassicaceae-insect interaction [14], within which intensive genomic and phenotypic information is available for a world-wide collection of natural accessions [15]. In *Arabidopsis*-herbivore interactions, plant trichomes (epidermal hairs) function as a mechanical barrier against feeding and oviposition by insect herbivores [11, 16, 17, 18]. Glucosinolates (GLSs) are major secondary metabolites of Brassicales that act as toxic chemicals against generalists [19, 20], but have the potential to attract specialist herbivores [14, 21]. For example, previous studies on *A. thaliana* focused on how these physical and chemical traits confer resistance against specific herbivore species, such as the small cabbage white butterfly *Pieris rapae* [22, 23], the diamond back moth *Plutella xylostella* [16, 20], and the green peach aphid *Myzus persicae* [24, 25]. However, knowledge remains limited about (i) how many insect species occupy *A. thaliana* in the field, (ii) whether plant defense traits contribute to insect abundance and community composition, and (iii) the host genes that are responsible for community members and overall community composition.

Laboratory environments are highly constant compared to naturally fluctuating environments; consequently, the phenotype in the laboratory might not be adequate to understand how genes function in the field [26, 27, 28]. The concept of using field studies to determine gene functions (coined *in natura* study: [26, 28]) is applicable to extended phenotypes, such as herbivore abundance and communities on plants. The molecular basis of anti-herbivore defense traits of *A. thaliana* has been studied in laboratory using natural accessions with respect to trichomes [29, 30, 31] and secondary metabolites [20, 32], and thus provides an ideal opportunity to distinguish the community-wide effects of single genes from naturally existing variation in particular defense traits [33]. For example, trichome density has heritable variation among natural accessions of *A. thaliana* [30, 31, 33]. Loss of function in a transcriptional factor gene, *GLABRA1* (*GL1*, also called *GLABROUS1*) results in glabrous phenotypes on leaf and stem surfaces in *A. thaliana* and related species [34, 35, 36, 38, 39] independent of root hair development [34, 40, 41]. Laboratory experiments have shown that the loss of function in *GL1* decreases resistance against leaf-chewing herbivores [22], and improves plant growth by saving the cost of defense [26]. However, these genetic effects remain unexplored in the field.

Natural accessions of *A. thaliana* possess various genetic backgrounds regarding their life-cycles in addition to defense traits [31, 42, 43]. In some geographical regions, a rapid life-cycle of *A. thaliana* allows themselves to accomplish two or more generations within a calendar year [42, 43, 44, 45]. The spring cohort of these accessions germinates, flowers, and produces seeds within spring and early summer. The summer cohort subsequently occurs that germinates and spends the summer season at a vegetative stage, and flowers and produce seeds during autumn [43, 44]. These life-cycles of *A. thaliana* accessions depend on the level of seed dormancy which can be attributed to allelic status in the *DELAY OF GERMINATION1* (*DOG1*) and *DOG6* gene [43, 45, 46], and the duration to flower development which is determined by *FRIGIDA*, *FLOWERING LOCUS C* and several other genes [42, 43, 47]. In wild populations of Europe, *A. thaliana* are attacked by herbivores from late spring to summer: generalist slugs and seed predators occur during late spring, and during summer various insects, such as beetles, moths, aphids, and aphidophagous parasitoids, occupy *A. thaliana* [19, 48, 49]. This seasonal schedule leads us to assume that the summer cohorts of *A. thaliana* should offer an ideal model system between wild *A. thaliana* and diverse herbivorous insects in the field.

Common garden experiments using single-gene mutants provide a powerful tool to determine the causal link between a particular gene and its phenotypes [38, 42, 50, 51]. In this study, we transplanted two glabrous mutants and 17 natural accessions of *A. thaliana,* for which trichome density and glucosinolate concentration vary across plants. In particular, we focused on *gl1-1* and *gl1-2* accessions, of which the former is a null trichome mutant derived from L*er*-1 accession and the latter is a hypomorphic mutant from Col-0 accession [41, 52]. In addition, the natural accessions were selected to cover variation in trichome density, GLSs content, and life-cycles [30, 32]. Common garden experiments with these plants were conducted in the two field sites, Switzerland and Japan, to identify common patterns between the two insect communities. Three specific questions were addressed: (i) is there heritable variation in herbivore abundance and community composition on each *A. thaliana* accession; (ii) which plant trait (physical, chemical, or other life-history traits) does influence herbivore abundance and community composition; (iii) does the loss of function of a single gene *GL1* alter insect abundance and community composition?

## Methods

### Plant materials and defense traits

*Arabidopsis thaliana* L., commonly known as thale cress or mouse-ear cress, is an annual weed native to Eurasia and naturalized in North America and East Asia [15]. This species is predominantly self-fertilizing [53] and, when plants are collected from the wild population or when mutants are isolated by mutagenesis, selfed seeds can be maintained as an inbred line called an “accession”. Weak dormancy and early-flowering accessions, such as Col-0 and L*er*-1 [45, 46], form both the spring and summer cohort owing to their rapid life-cycle [43]. The spring cohort flowers and seed sets in spring, and then the summer cohort germinates in early summer and flowers in autumn [43]. The accessions with strong dormancy, such as Cvi-0 and Shahdara [46], pass summer as seeds, and furthermore the accession with strong dormancy and late-flowering phenotype, such as Kas-2, are predominantly a winter-annual that has only one generation within a calendar year [43]. In the wild populations, generalist slugs and seed weevils feed on *A. thaliana* during late spring, while more diverse herbivores, such as *Phyllotreta* beetles, green peach aphids *Myzus persicae*, and diamondback moths *Plutella xylostella*, occur during summer [48, 49]. The summer cohort of *A. thaliana* remains at the vegetative stage during summer [43] and thereby provides various herbivores with an opportunity to feed on vegetative plants in the field. We transplanted vegetative *A. thaliana* during June and July to the field sites to simulate the summer cohorts (see ‘Common garden experiment’ section for details).

To cover wide variation in trichome density (physical defense) and GLS accumulation (chemical defense) with early- and late-flowering cycles, we selected 17 natural accessions and two glabrous mutants (Table 1). The natural accessions selected in this study should represent the world-wide genetic variation, because the genome-wide pairwise genetic distance was 5.7% in median, which is comparable to that of all accessions analyzed by the 1001 Genome Consortium [15]. These 17 accessions include both early- and late-flowering accessions (e.g. Col-0 and Kas-2 analyzed by Taylor et al. [43]), such that the flowering time under a long-day laboratory condition ranges from 23 (Ws-2 accession) to 92 days (Br-0 accession) [31]. To examine the effects of plant life-history traits on insect community composition, we measured and incorporated the plant size and presence of flowering stem (see ‘Common garden experiment’ and ‘Statistical analysis’ below).

**Table 1.**
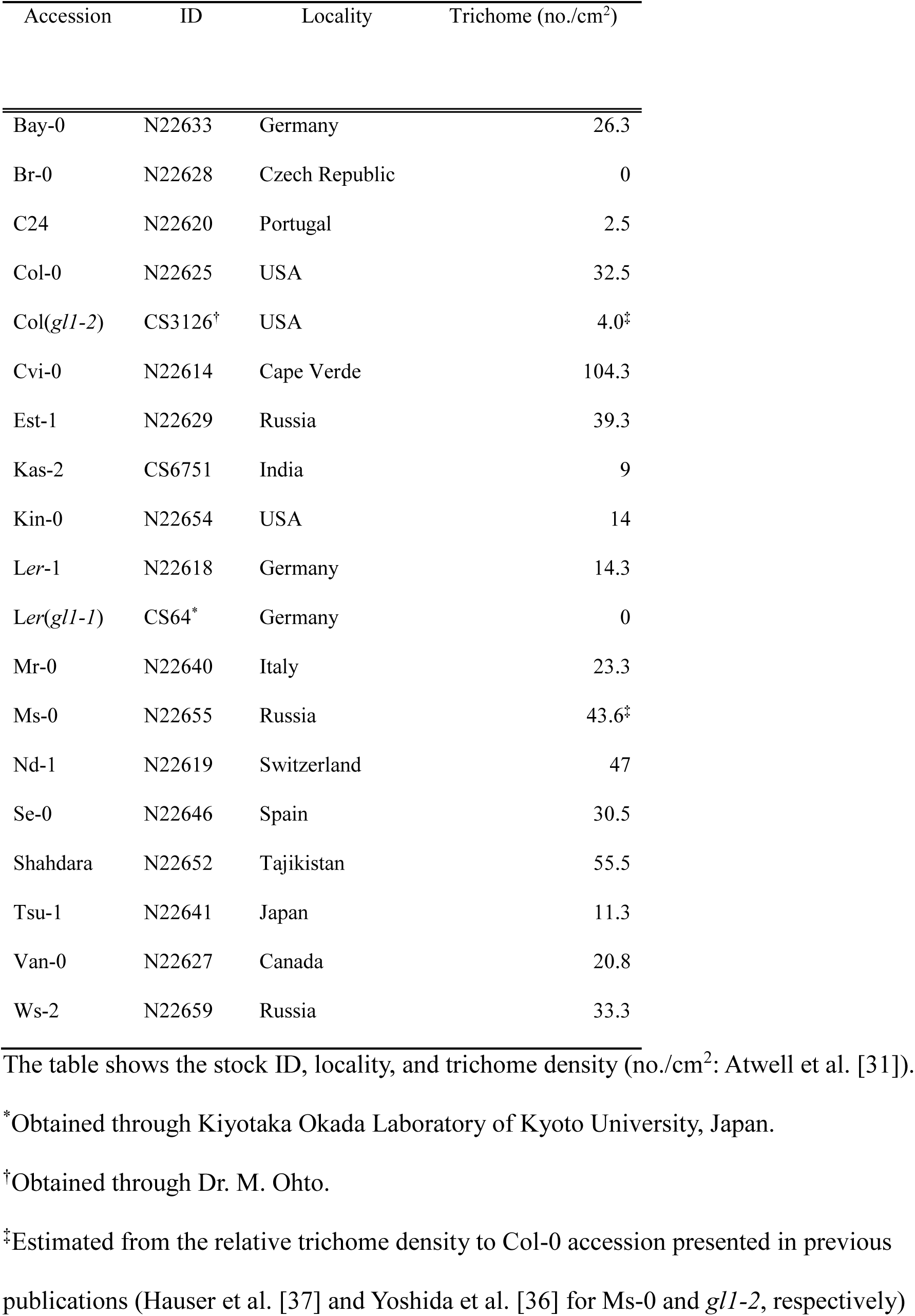
*Arabidopsis thaliana* accessions used in this study.

To test the functional advantage of *GL1* gene in producing trichomes, we added two glabrous mutants, *gl1-1* and *gl1-2*, to the set of natural accessions (Table 1). The former mutant *gl1-1* has the background of L*er* accession with null mutation due to a 6.5-kb deletion on *GL1* and lacks leaf surface trichomes. The latter *gl1-2* has the background of Col accession with the deletion of 27 amino acid induced by X-ray radiation, showing hypomorphic mutation with lower density of trichome on leaf surface [40, 52]. Out of the 17 natural accessions, Br-0 and C24 have no or few trichomes due to a frameshift mutation and one amino acid change in the myb DNA binding domain of GL1, respectively [30]. We compiled the data of leaf trichome density (no./cm^2^) from the GWA-portal (https://gwas.gmi.oeaw.ac.at/: [37]).

All natural accessions were included in previous quantitative genetic studies of GLS, of which seven accessions were used as parental genotypes of recombinant inbred lines (e.g., Col × L*er* and Cvi × L*er* [20]; Bay × Sha [54]; Kas × Tsu [55]) and the other accessions were used in a genome-wide association mapping [34]. To test whether genetic potentials in GLS profiles explain herbivory rate, we used the data in Chan et al. [32] on 21 GLSs of 96 *A. thaliana* accessions using a mature leaf at 35-days post germination from a plant grown under short-day laboratory conditions without herbivory. As they performed two trials to quantify GLS, we used the average GLS contents (nmol/mg flesh weight). We focused on variation in aliphatic GLSs and its chain-length, because these parameters play a major role in preventing above-ground herbivory [50, 56, 57]. Regarding the data of Chan et al. [32], we applied a principal component analysis (PCA) to Total C3-, C4-, C5-, C7-, and C8-aliphatic GLSs. The first and second principal components explained 44% and 33% variation in the GLS profiles among our 17 accessions, respectively (Fig. S1); therefore, these two components were used in our statistical analyses.

### Common garden experiment

We used the experimental gardens of the University of Zurich at Irchel campus (Zurich, Switzerland: 47° 23′ N, 8° 33′ E, alt. ca. 500 m) and the Center for Ecological Research, Kyoto University (Otsu, Japan: 35° 06′ N, 134° 56′ E, alt. ca. 200 m) (Fig. 1). The Zurich site is close to a deciduous forest and the surroundings of the common garden are covered with concrete tiles to prevent weeds. The Otsu site is a suburb of cultivated fields and the ground of the study site is covered with short grasses. In the Otsu site, the grass weeds were mown and the surroundings were covered with agricultural sheets before the experiment. At both sites, no large *Brassica* plants occur during early summer. Average air temperature and total precipitation was 19 °C and 198 mm in Zurich (during July 2016; MeteoSwiss, http://www.meteoswiss.admin.ch/home.html) and 22 °C and 321 mm in Otsu (during June 2016; Japan Meteorological Agency, http://www.jma.go.jp/jma/index.html).

**Figure 1.**
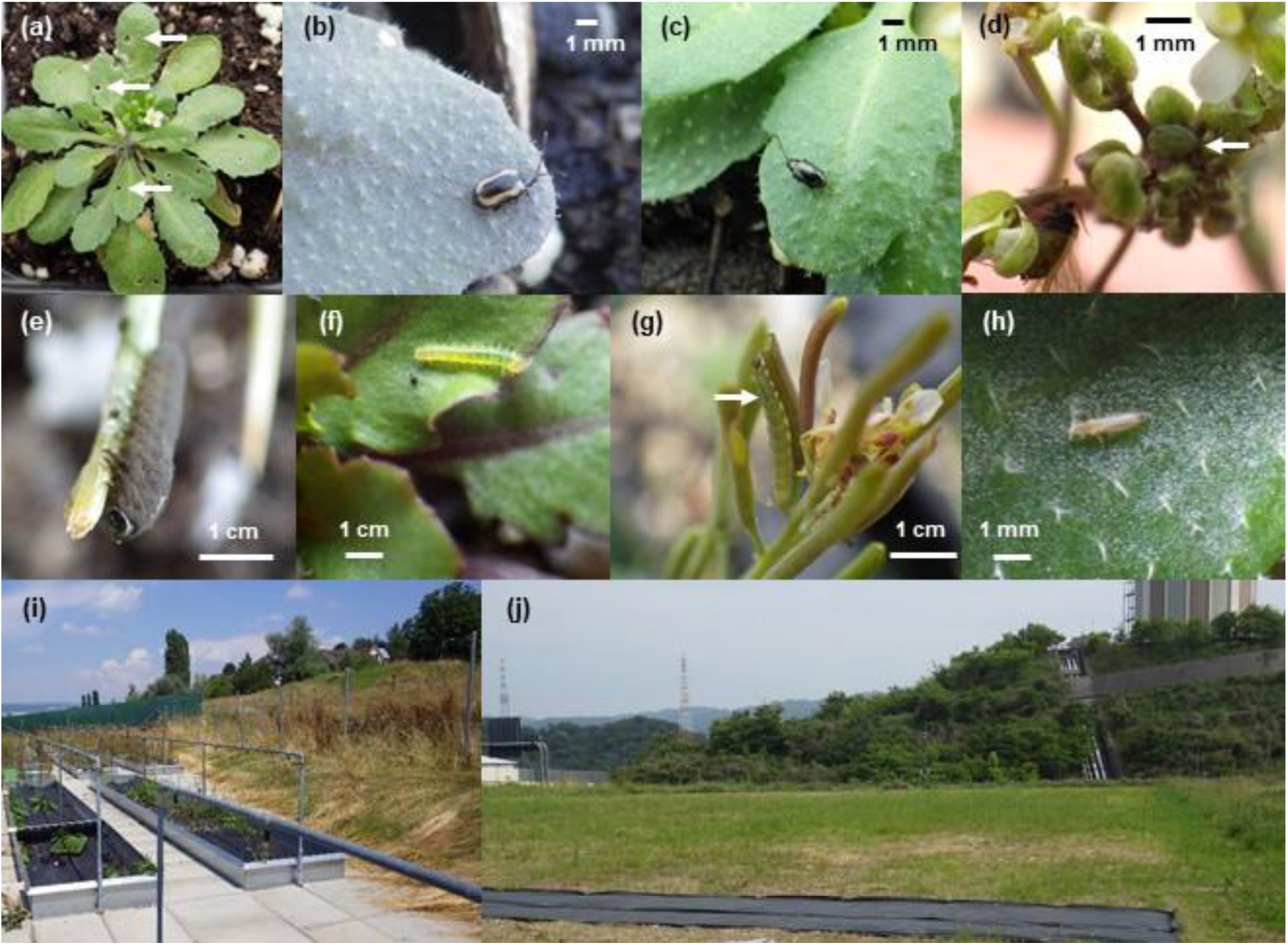
Photographs of plants and insects. **(a)** Leaf holes made by flea beetles (arrows), **(b)** a striped flea beetle *Phyllotreta striolata*, **(c)** a turnip flea beetle *Phyllotreta atra*, **(d)** mustard aphids *Lipaphis erysimi*, **(e)** a larva of the turnip sawfly *Athalia rosae*, **(f)** a newly hatched larva of the small cabbage white butterfly *Pieris rapae* **(g)** a larva of the diamond back moth *Plutella xylostella*, **(h)** a western flower thrips *Frankliniella occidentalis*, **(i)** the field site in Zurich, Switzerland, and **(j)** the field site in Otsu, Japan.

We prepared 10 replicates of 19 accessions (= 190 plants in total) for each experiment. Experimental plants were initially grown in an environmental chamber, and were then transferred to the outside garden. To cultivate plants, we used mixed soils of agricultural composts (Profi Substrat Classic CL ED73, Einheitserde Co. in Switzerland; MetroMix 350, SunGro Co. in Japan) and perlites with a compost to perlite ratio of 3:1 litter volume. No additional fertilizers were supplied because the agricultural soils contain fertilizers. Seeds were sown on the soil and stratified under constant dark conditions at 4–5 °C air temperature for a week. Plants were then grown under short-day condition (8h:16h light:dark [L:D], 20 °C air temperature, and 60% relative humidity) for 1 month to prevent flowering before the field experiment. The plant positions were rotated every week to minimize the growth bias by light condition. Plant individual was moved to a plastic pot (7.5-cm diameter, with 6.5-cm depth in Japan; 6.0 × 6.0 × 6.0 cm in Switzerland), and acclimated for 3 days at shaded outdoor place before the field experiments. The potted plants were randomly placed among three blocks in each common garden: 68, 69, and 53 plants were assigned within each block in Zurich; and 76, 76 and 38 plants were assigned within each block in Otsu. The potted plants were set in a checkered manner within a block without being embedded in the ground on water-permeable plastic sheet. Each block was more than 1.0 m apart from each other. These experiments were conducted from June 18 to July 1, 2016 in Otsu; and from July 13 to August 3, 2016 in Zurich. Plants were watered every three days in Otsu and every day in Zurich.

Insect and herbivorous collembola on individual plants were visually counted every 2–3 days. These species were identified ocularly with a magnifying glass. Dwelling traces and mummified aphids were also counted as a proxy of the number of leaf miners and parasitoid wasps, respectively. Eggs, larvae, and adults were counted for all species, as long as they could be observed by the naked eye. The abundance of each species was evaluated by the cumulative number of individuals over the experimental period to reflect herbivory load on plants [58]. Small holes made by flea beetles were counted at the Zurich site and the maximum number throughout the experiment was used as an indicator of damage by flea beetles; however, this phenotyping was difficult in Japan, due to heavier and simultaneous infestation by sawflies. We attempted, but failed, to evaluate leaf damage in Japan because about one third of individual plants were dead by the end of the experiment due to high air temperature in June. All counting was conducted by a single observer during the daytime (08:00–17:00), and was continued for 3 weeks after the beginning of the field experiment.

We recorded the initial plant size and presence/absence of flowering stems to incorporate the effects of plant life-history traits on insect abundance. Initial plant size was evaluated by the length of the largest rosette leaf (mm) at the beginning of the field experiment, because this parameter represents plant size at the growth stage. The presence/absence of flowering stems was recorded 2 weeks after transplanting plants.

### Statistical analysis

*Response variables -* Community indices were examined at three levels (i.e., component species, guilds, and entire communities) as response variables in the following analyses. At the species level, we analyzed the number of individuals of each herbivorous species. We analyzed species for which more than 20 individuals were observed in each site, because statistical tests were difficult to apply to rare species. For the Zurich data, we analyzed the number of leaf holes as an indicator of damage by flea beetles. At the guild level, we classified herbivorous species into those feeding on external leaf tissues (i.e., leaf chewers) and those feeding on internal plant tissues (including sap suckers and leaf miners). We also separated herbivorous species into specialists on Brassicaceae (e.g., white butterflies, cabbage sawflies, and turnip flea beetles) and generalists on multiple plant families (some species of aphids and thrips) (Table 2; Fig. S2). The total number of insect individuals in each category was analyzed as guild level statistics. At the entire community level, we calculated species richness (i.e., number of species), Shannon’s diversity index *Hʹ*, and the total number of insect individuals on individual plants. All of the response variables were ln(*x* + 1)-transformed to improve a normality before statistical analyses. All statistical analyses were conducted using R version 3.2.0 [59]. We utilized the *rda* function (in the *vegan* package: [60]) to perform the redundancy analysis. We used the *lme* function (in the *nlme* package: [61]) to estimate heritability, as described below.

**Table 2.** Insect species observed on field-grown *Arabidopsis thaliana*.

| Common name | Scientific name | Feeding habit | Host range | Abundance <sup>†</sup> |  |
| --- | --- | --- | --- | --- | --- |
|  |  |  |  | Zurich | Otsu |
| Cabbage looper | *Trichoplusia ni* | Leaf chewer | Generalist | 0 | - |
| Diamond back moth | *Plutella xylostella* | Leaf chewer | Specialist | ++ | ++ |
| Garden springtail* | *Bourletiella hortensis* | Leaf chewer | Generalist | - | + |
| Piggyback grasshopper | *Atractomorpha lata* | Leaf chewer | Generalist | - | - |
| Small cabbage white butterfly | *Pieris rapae* | Leaf chewer | Specialist | + | + |
| Striped flea beetle | *Phyllotreta striolata* | Leaf chewer | Specialist | ++ | - |
| Turnip flea beetle | *Phyllotreta atra* | Leaf chewer | Specialist | +++ | 0 |
| (Leaf holes made by flea beetles) | *Phyllotreta* spp. | Leaf chewer | Specialist | +++ | - |
| Turnip sawfly | *Athalia rosae* | Leaf chewer | Specialist | 0 | +++ |
| (Dwelling traces) | NA | Internal feeder | Generalist | - | 0 |
| Green peach aphid | *Myzus persicae* | Internal feeder | Generalist | + | + |
| Mustard aphids | *Lipaphis erysimi* | Internal feeder | Specialist | +++ | + |
| Onion thrip | *Thrips tabaci* | Internal feeder | Generalist | + | + |
| Western flower thrip | *Frankliniella occidentalis* | Internal feeder | Generalist | ++ | ++ |
| (Parasitoid wasp indicated by mummified aphids) | NA | Carnivore |  | + | + |
| Seven-spot ladybird | *Coccinella septempunctata* | Carnivore |  | 0 | - |

Detailed abundance in the Zurich and Otsu sites is provided in the supplementary material

*Variation in insects on plant accessions* - To quantify variation in insect communities among plant accessions and study sites, we performed a redundancy analysis to partition sources of variation in community composition into the plant accession, study sites, and accession-by-site effects. The accession-by-site interaction was first analyzed by 999-times permutation tests, and then the main effects of accessions and sites were examined without the interaction term. Then, we estimated broad-sense heritability *H*^2^ in a focal response, as the proportion of variance attributable to plant accessions. We used liner mixed models, in which the accession ID was assigned as a random effect. This variance component of random effect was estimated by the restricted maximum likelihood method [62, 63]. The significance of heritability was examined by likelihood ratio tests by comparing the linear models with or without the random effect of accession ID. This estimation of heritability was separately performed for the data from Zurich and Otsu. *P*-values were corrected by the false discovery rate (FDR) of multiple testing [64]. Another option of estimating heritability is to incorporate a genetic distance matrix among natural accessions, as used in genome-wide association studies [e.g. 65, 66]. However, it was difficult to apply the same approach to single-gene mutants and the limited number of accessions; thus, we adopted the linear mixed models without the distance matrix to estimate broad-sense heritability.

*Effects of plant traits* - To address whether particular plant traits contributed to community members and composition, we used multiple regressions that considered trichome density, PC1 and PC2 of aliphatic GLSs, the presence/absence of flowering stems, and initial plant size (mm) as explanatory variables. No explanatory variables were heavily correlated with each other (|*r*| < 0.6 for all pairs). We considered the difference of experimental block as a covariate. Because trichome density had a highly skewed distribution, due to completely glabrous phenotypes, this variable was ln(*x* + 1)-transformed before the analysis. First, we tested the effects of plant traits on each response variable without the dataset on glabrous mutants. When detecting significant effects of trichomes on a particular herbivore, we then compared two glabrous mutants and their parental accessions to test how the *GL1* genes impact guild and community indices encompassing the focal herbivore. Linear mixed models were used to analyze trichome production, initial plant size (mm), and the presence/absence of flowering stems as explanatory variables. The difference in parental background (i.e., L*er*-1 or Col-0) was considered as a random effect. We used the *lme* function with the maximum likelihood method for these mixed models. All of the continuous response and explanatory variables were standardized following a normal distribution, with zero mean and one variance, to make coefficients comparable between the linear models. *P*-values were corrected by FDR [64].

## Results

### Abundance and communities of insects among plant accessions and study sites

We observed 15 insect species including flea beetles, sawflies, butterflies, moths, aphids, and thrips on *A. thaliana* in the two field experiments (Table 2; Fig. 1). Of these insects, 5 and 3 species were specific to the Otsu and Zurich site, respectively. Redundancy analysis and permutation tests confirmed that the plant accession, study site, and accession-by-site effects exhibited significant sources of variation in the community composition (Accession, Sum of Squares (SS) = 0.99, *F* = 1.57, *P* < 0.001; Site, SS = 1.31, *F* = 36.2, *P* < 0.001; Accession-by-site, SS = 0.96, *F* = 1.54, *P* < 0.001 with 999 permutations; Fig. 2). We found significant broad-sense heritability in species richness, Shannon diversity, and total abundance of insects on *A. thaliana*, and its magnitude varied between two study sites (10-11% and 15–30% heritability in Zurich and Otsu, respectively: Table 3). When each community member was analyzed separately, we found significant 16% and 33% heritability in the abundance of two predominant herbivores, the striped flea beetle *P. striolata* in Zurich and the turnip sawfly *A. rosae* in Otsu, respectively (Table 3). We detected 40% heritability in the number of leaf holes made by flea beetles in Zurich (Table 3). At both sites, significant heritability was detected in the number of herbivorous individuals for each of the leaf chewer, specialist, and generalist guilds rather than in single species (Table 3). Even when the two mutants *gl1-1* and *gl1-2* were eliminated from our dataset, heritability remained at a similar level with respect to the abundance of the two predominant herbivore species (striped flea beetle in Zurich, *H*^2^ = 0.20, 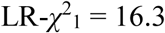, *P*_fdr_ < 0.001; turnip sawfly in Otsu, *H*^2^ = 0.31, 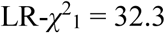, *P*_fdr_ < 10^−6^), the abundance of leaf-chewing herbivores (Zurich, *H*^2^ = 0.13, 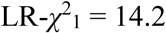, *P*_fdr_ < 0.001; Otsu, *H*^2^ = 0.28, 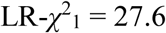, *P*_fdr_ < 10^−6^), and insect species richness (Zurich, *H*^2^ = 0.14, 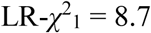, *P*_fdr_ < 0.01; Otsu, *H*^2^ = 0.28, 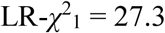, *P*_fdr_ < 10^−6^).

**Figure 2.**
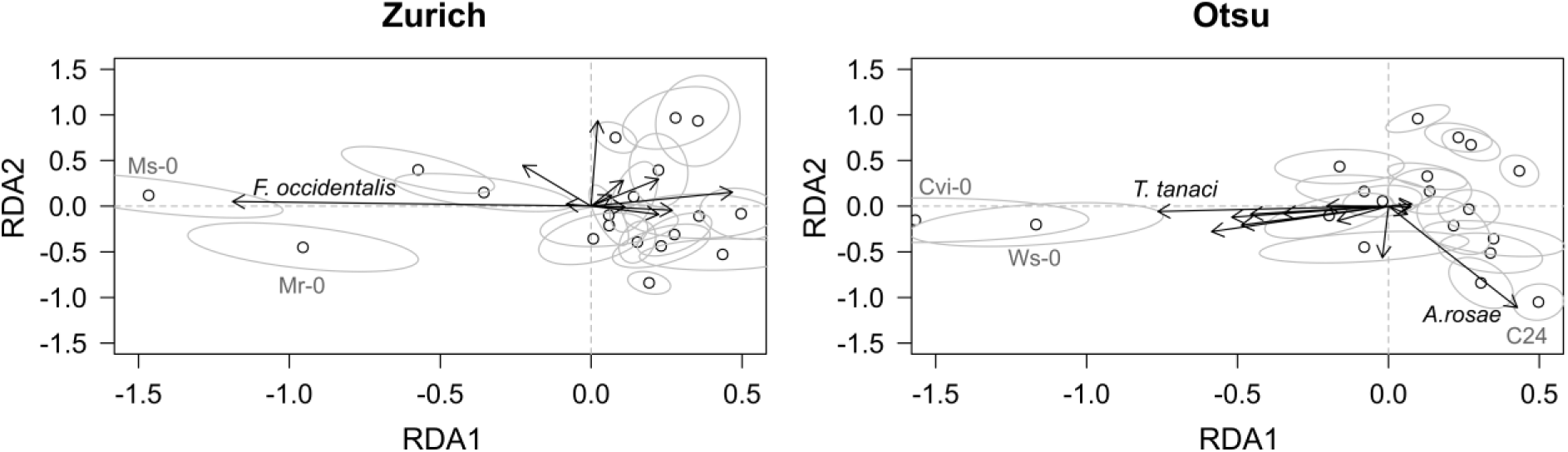
Redundancy analysis summarizing the community composition among 19 accessions of *A. thaliana* in Zurich, Switzerland and Otsu, Japan. White and grey circles indicate the accession mean and its standard error projected on the first and second RDA dimension. Arrows represent the contributions of each species. Permutation tests confirmed significant variation in the community composition among plant accessions and study sites (see the Results section).

**Table 3.** Likelihood ratio tests for estimating broad-sense heritability *H*^2^.

| Response | Zurich |  |  | Otsu |  |  |
| --- | --- | --- | --- | --- | --- | --- |
| | $H^2$ | LR-$\chi^2$ | $P_{\text{fdr}}$ | $H^2$ | LR-$\chi^2$ | $P_{\text{fdr}}$ |
| Turnip sawfly | --- | --- | --- | **0.33** | **41.0** | **2.0.E-09** |
| Leaf holes | **0.40** | **55.1** | **1.5.E-12** | --- | --- | --- |
| Striped flea beetle | **0.16** | **12.6** | **1.2.E-03** | --- | --- | --- |
| Turnip flea beetle | 0.01 | 0.16 | 0.75 | --- | --- | --- |
| Diamond back moth | 1.6E-09 | -6.5E-08 | 1 | 0.07 | 2.87 | 0.13 |
| Small cabbage white butterfly | --- | --- | --- | 0.05 | 1.92 | 0.22 |
| Green peach aphid | --- | --- | --- | 0.03 | 0.55 | 0.50 |
| Mustard aphid | 0.04 | 1.30 | 0.32 | 0.03 | 0.48 | 0.50 |
| Western flower thrip | **0.32** | **38.8** | **3.1.E-09** | 0.03 | 0.46 | 0.50 |
| Leaf chewer | **0.14** | **10.4** | **0.003** | **0.28** | **31.3** | **9.4.E-08** |
| Internal feeder | 0.04 | 1.22 | 0.32 | **0.14** | **9.7** | **0.003** |
| Specialist | **0.11** | **6.32** | **0.02** | **0.25** | **25.8** | **1.0.E-06** |
| Generalist | **0.17** | **13.2** | **0.001** | **0.12** | **7.5** | **0.01** |
| Species richness | **0.11** | **6.82** | **0.02** | **0.26** | **27.3** | **5.8.E-07** |
| Shannon diversity | **0.11** | **6.10** | **0.02** | **0.17** | **13.4** | **0.001** |
| Total abundance | **0.10** | **5.97** | **0.02** | **0.29** | **33.0** | **5.9.E-08** |

Likelihood ratio, LR- $\chi^2$ , was tested by comparing the models with and without a random effect of the accession ID. $P$ -values were based on a $\chi^2$ distribution with one degree of freedom and corrected by false discovery rate, FDR [64]. Bold values indicate significant $H^2$ at $P_{\text{fdr}} < 0.05$ . Bars indicate no information available due to low abundance.

### Plant traits underlying the abundance and communities of insects

We examined whether the species, guild, and community structure of insects were affected by trichome density, glucosinolates, and life-history traits among natural accessions (Fig. 3). Significant effects of trichomes at the species and guild levels were observed (Fig. 3; Table S1). Two predominant leaf chewers, the striped flea beetle in Zurich and the turnip sawfly in Otsu, occurred less on hairy accessions than on accessions that produced low quantities of trichomes (Fig. 3, 4b; Table S1). The number of leaf holes was smaller on hairy plants compared to glabrous plants at the Zurich site (Fig. 3, 4a; Table S1), indicating that trichomes have a resistance function against flea beetles. At the Otsu site, the abundance of the eggs and larvae of the small cabbage white butterfly *Pieris rapae* was also low on hairy plants (Fig. 3; Table S1). At the guild level, trichomes had significant negative effects on the leaf chewers at both sites (Fig. 3, 4c, 4d; Table S1). In contrast to trichomes, aliphatic GLSs did not have any consistent effects on herbivore abundance. The first principal component of GLSs had negative effects on leaf chewers, specialist herbivores, species richness, and total abundance at the Zurich site but no significant effects on these indices at the Otsu site (Fig. 3). The second principal component of GLSs was positively correlated with the abundance of turnip sawfly at the Otsu site and western flower thrips at the Zurich site (Fig. 3: Table S1). These effects of trichomes and GLSs were variable between the two sites with respect to insect richness, Shannon diversity, and total abundance (Fig. 3, 4e, 4f; Table S1). Initial plant size or the presence of flowering stems significantly increased insect richness, diversity, and total abundance at the both sites (Fig. 3; Table S1). The result that Kas-2 in Otsu and C24 in Zurich were less likely occupied by leaf chewers (Fig. 4) was due to its small plant size.

**Figure 3.**
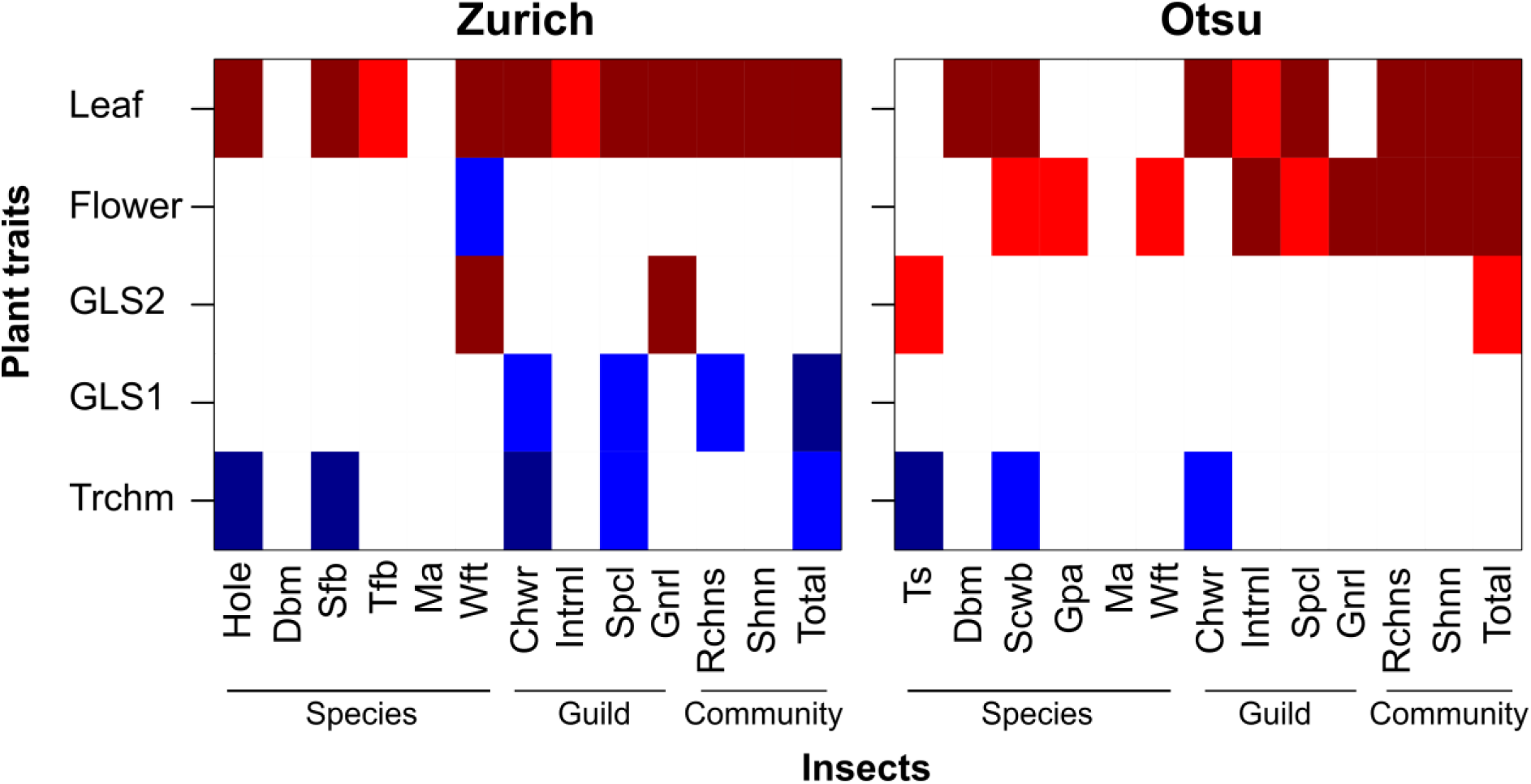
A heat map showing FDR-corrected p-values (*P*_fdr_) for the effects of plant traits on insect species, guild, and community indices among 17 natural accessions in Zurich, Switzerland and Otsu, Japan. Shown are the effects of trichome density (Trchm), PC1 and PC2 of aliphatic glucosinolates (GLS1 and GLS2), presence of flowering stems (Flower), and initial leaf length (Leaf) on the diamond back moth (Dbm), striped flea beetle (Sfb), turnip flea beetle (Tfb), mustard aphid (Ma), western flower thrip (Wft), turnip sawfly (Ts), small cabbage white butterfly (Scw), green peach aphid (Gpa), leaf chewers (Chwr), internal feeders (Intrnl), specialists (Spcl), generalists (Gnrl), species richness (Rchns), Shannon diversity (Shnn), and total abundance (Total). Colors represent the sign and significance of trait effects: 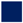(dark blue), - coef. with *P*_fdr_ < 0.01; 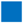(blue), - coef. with *P*_fdr_ < 0.05; 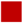(dark red), + coef. with *P*_fdr_ < 0.01; 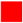(red), + coef. with *P*_fdr_ < 0.05; 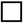(white), not significant at *P*_fdr_ > 0.05.

**Figure 4.**
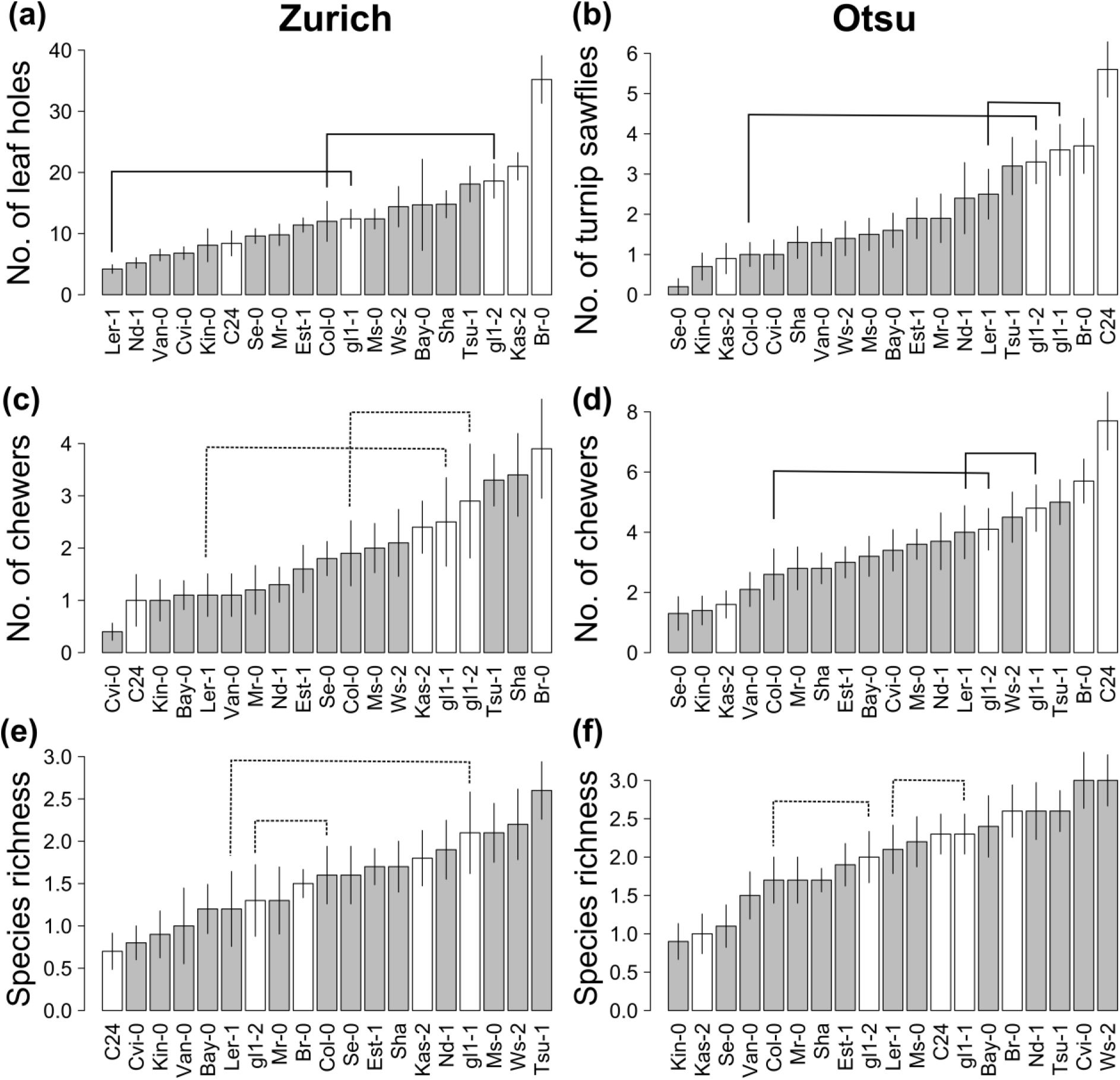
Variation in insect species, guild, and community on 19 *A. thaliana* accessions (mean ± SE) in Zurich, Switzerland (left panels) and Otsu, Japan (right panels). Given the significant effects of trichomes on flea beetles *P. striolata* and sawflies *A. rosae* (Fig. 3), these panels show herbivory, guild, and community indices comprising the flea beetles and sawflies. White bars of plant accessions represent the sparse density of less than 10 leaf trichomes/cm^2^. Connected lines highlight pairs between a glabrous mutant and its parental accession, where solid and dashed lines indicate significant and non-significant differences between the mutants and parental accessions at *P*_fdr_ < 0.05.

### Comparing glabrous mutants and parental hairy accessions

We examined the effects of a single gene *GLABRA1* on herbivory, guild, and community indices encompassing two predominant leaf chewers, flea beetles and sawflies. At the species level, compared to parental accessions, two glabrous mutants had significantly more leaf holes made by the flea beetles and larvae of the turnip sawfly (

Fig. 4a, 4b; Table S2). At the guild level, leaf chewers tended to occur more often on two glabrous mutants than on each of the parental accessions (Fig. 4c, 4d), although this difference was not statistically significant in Zurich (Table S2). Among community indices, total abundance at the Otsu site was significantly lower on glabrous mutants than on hairy parents (Fig. 4e, 4f) (coef. ± SE = −0.46 ± 0.16, *Z* = −2.86, *P* < 0.01; Table S2).

## Discussion

*Arabidopsis thaliana* is the best-studied plant in laboratories, and is distributed across the temperate region [15]. Several researchers have studied leaf herbivory [17, 50, 56], herbivorous fauna [19, 48, 49, 67], and plant fitness [17, 48, 50, 56] in *A. thaliana* under field conditions; however, quantitative evidence remains limited in relation to insect community composition on this plant species. In the present study, we found that the community composition of 15 arthropod species was a significantly heritable phenotype for *A. thaliana* in the two distant field sites. Importantly, the loss of function of the *GLABRA1* (*GL1*) gene significantly decreased plant resistance against two predominant chewers, flea beetles and sawflies, at the species level. At the guild level, both *gl1-1* and *gl1-2* mutant plants were more likely to be inhabited by leaf chewers than each of their hairy parents L*er*-1 and Col-0 at both the Zurich and Otsu sites (Fig. 1c, 1d). The parallel pattern in *gl1-1* and *gl1-2* suggests that the single gene *GL1* contributes to community composition. These results demonstrate that variation in a single gene contributes towards shaping insect communities through its impacts on leaf-chewing herbivores.

### Heritable variation in insect community composition

Despite the large difference in insect community composition, we consistently found significant heritability and effects of trichomes in the two field sites. Our estimated heritability seems moderate (Table 3), but is comparable with the other studies reporting less than 50% heritability in insect abundance and community composition [6, 7, 8]. Using plant genotypes propagated from seeds, Johnson and Agrawal [6] showed that heritability in insect species richness on the perennial *Oenothera biennis* ranged from 10 to 40%, depending on habitat conditions. Our present study also used *A. thaliana* accessions with unique genetic backgrounds, resulting in the moderate heritability of species richness and diversity. We also found that the presence of flowering stems or larger plant size increased species richness and diversity (Fig. 3; Table S1). Plant apparency hypothesis posed the importance of such plant life-history traits in anti-herbivore defense [68], and this hypothesis has been recently supported by a meta-analysis [69], comparative study [70], and genome-wide association mapping [65]. In the context of community genetics, plant life-history traits are a key predictor of insect community composition on perennial herbs [6] and woody plants [2]. Our present study supports the plant apparency hypothesis at the community level of insects on *A. thaliana*.

Consistent with our results in the Zurich site, Harvey et al. [49] found that *A. thaliana* plants sown in summer were heavily attacked by *Phyllotreta* beetles and also harbored by diamondback moths and aphids in the Netherlands. By simulating a summer cohort of *A. thaliana* with various background accessions of flowering time and seed dormancy, our present study showed the importance of plant life-history traits in organizing the summer insect community. In the seasonal context, the parental accessions of two glabrous mutants, Col-0 and L*er*-1, have a weak dormancy allele of *DELAY OF GERMINATION* (*DOG1*) [46], which allows *A. thaliana* to germinate during summer [43, 45] and thus to serve as food plants for summer herbivores in the field. As we have shown the defense advantage of producing leaf trichomes, the present field experiment could represent how the loss of function in *GL1* imposes summer herbivory on weak dormancy accessions. The seasonality of plant defense would motivate us to further study epistasis or pleiotropy among genes involved in antiherbivore defense and seasonal phenology of *A. thaliana*.

### Effects of the single gene GLABRA1 on insect abundance

In reverse genetic analysis, multiple independent mutants with a consistent phenotype are required to prove the function of a particular gene. In addition, using multiple genetic backgrounds of parent can also give a strong proof of a gene function. In a previous study, the roles of single genes related to GLS biosynthesis in modulating herbivory were quantified using mutants derived from a single parental accession, Col-0 [50]. Our common garden experiments illustrate the function of *GL1* gene against herbivory *in natura* using two distinct lines, L*er* (*gl1-1*) and Col (*gl1-2*). To date, several studies have reported associations between *GL1* polymorphism and anti-herbivore functions in field populations of *A. lyrata* [9, 10] and *A. halleri* [11, 18]. Plant trichomes also prevent herbivory by sawflies [71] and flea beetles [72, 73] on *Brassica* cultivars. Together with these results, our present results indicate that plant trichomes and a single gene *GL1* play a key role in physical defense against crucifer feeders.

Laboratory experiments on single-gene mutants and natural accessions of *A. thaliana* suggested that plants with high trichome density resisted infestation by aphids [25, 74]. Under the two tested field conditions, trichomes had no significant effect on the abundance of aphids, possibly because aphids primarily occurred on flowering stems, on which the trichome density is low. In fact, the presence of flowering stems was positively correlated with the abundance of aphids (Table S1). These results support the limited associations between aphid abundance and *GL1* polymorphism detected in field-grown *A. halleri* [11]. In addition, we could not detect any significant effects of trichomes and *GL1* on the abundance of larval *P. xylostella*, even though trichomes prevent adult moths ovipositing on *A. thaliana* under laboratory conditions [16]. Handley et al. [16] focused on several northern accessions of *A. thaliana*, whereas the current experimental setting covers a wider geographical range of natural accessions. A recent genome-wide association study using 350 natural accessions also found no significant association between *GL1* polymorphism and herbivory by *P. xylostella* [66]. Combined with the previous studies, our present results from field-grown *Arabidopsis* exemplify the importance of testing diverse accessions and environmental conditions.

Although several studies on *Nicotiana* plants illustrated the effects of single jasmonate signaling genes on herbivore abundance and communities, silencing jasmonate pathway results in complex pleiotropy on multiple defense traits in leaves and flowers [12, 13, 75, 76]. In contrast, *Arabidopsis* trichomes have a simple molecular mechanism that allows *GL1* to be a prime candidate gene for physical defense without pleiotropy. Loss of function mutants in a few transcriptional factor genes (including *GLABRA1* (*GL1*), *GLABRA2*, *GLABRA3*, *TRANSPARENT TESTA GLABRA1*) result in glabrous phenotypes in *A. thaliana*. While the loss of function of the latter three genes results in pleiotropic defects in root hairs, the loss of function of *GL1* does not affect root hairs, due to the subfunctionalization of *GL1* and its homolog *WEREWOLF* [34, 35, 41, 52]. Indeed, many independent null or hypomorphic mutations of *GL1* have been reported in natural accessions of *A. thaliana* [30, 37]. Of note, the Br-0 and C24 accessions were the most susceptible accessions to leaf chewers at each site (Fig. 4c, 4d), and have disruptive mutations on *GL1* [30, 37]. Based on genetic regulatory systems and natural variation, the present findings on *GL1* confer an evolutionary implication to its functional advantage in producing trichomes against herbivory.

### Varying effects of chemical defense on specialist herbivores

Glucosinolates act as a chemical defense against herbivory [19, 20, 56, 57]; however, some specialist herbivores overcome GLSs [20, 57, 77, 78]. Because the insect communities observed here were mainly composed of specialist herbivores (Table 2), aliphatic GLSs might have had variable effects in our present study. Specifically, the striped flea beetle *P. striolata* efficiently sequesters 4-methylthiobutyl from *A. thaliana*, a short-chain aliphatic GLS [78]. The larvae of *A. rosae* sawflies also sequester GLSs [77], whereas adults utilize isothiocyanates, which are breakdown products of aliphatic GLSs, to find host plants [21]. The sequestration and host-finding might explain the result that some components of GLSs had negative effects whereas the others had no or positive effects on the abundance of specialist herbivores.

The present study utilized GLS data quantified under laboratory conditions with an aim to address whether genetically based variation in GLSs is associated with insect communities. However, these genetic potentials might be insufficient to reflect the effects of GLSs on herbivore abundance in the field, due to phenotypic plasticity and induced response of GLSs to herbivory [23, 24, 50]. For example, the green peach aphid *M. persicae* and the cabbage white butterfly *P. rapae* can modify the expression level of the *MAM1* gene, which involves a chain elongation of aliphatic GLS [23, 24]. In a field study, Kerwin et al. [50] found gene-by-environmental effects on GLS profiles and herbivory on *A. thaliana*, and these effects varied considerably among study years and sites. Multi-year surveys are therefore needed to reveal under what conditions GLS profiles contribute to shaping insect community composition.

### Conclusion

Our field investigation showed a genetic basis in the insect community assemblage on *A. thaliana*, and the advantage of the functional allele of *GL1* in avoiding leaf chewers. In Brassicaceae plants, evidence is accumulating to suggest that genetic variation within a plant species alters insect community composition and, in turn, exerts selection on plant defense [4, 79, 80]. Variation in the trichome density is also observed across Brassicaceae plants [71, 72, 73], where *GL1* orthologs affect the trichome density [81]. In the context of community genetics, the present study on *GL1* provides evidence of a key gene affecting the community composition of crucifer-feeding insects. Future study should assess the relative importance of single genes and quantitative genetic variation towards a complete understanding of plant genetic effects on insect community assembly.

## Abbreviation

*GL1*: *GLABRA1*; GLS: glucosinolate; PCA: principal component analysis; RDA: redundancy analysis; FDR: false discovery rate

## Acknowledgements

The authors thank Jordi Bascompte and Matthew Barbour for their valuable discussions, and Satoshi Oda and Aki Morishima for helping with plant cultivation and fieldwork.

## Funding

This study was supported by JSPS Postdoctoral Fellowship (Grant Number, 16J30005) and JST PRESTO (JPMJPR17Q4) to Y.S., JST CREST (JPMJCR15O2) and JSPS KAKENHI (JP16H06171 and JP16H01473) to A.J.N., Swiss National Foundation, URPP Global Change and Biodiversity of the University of Zurich, JST CREST (JPMJCR16O3), Japan to K.K.S. The field experiment in Japan was supported by the Joint Usage/Research Grant of Center for Ecological Research, Kyoto University.

## Availability of data and source codes

The data and R source code are included in the online supplementary material (Additional_file1_Data.xls; Additional_file2_Rscript.txt).

## Authors’ contributions

Y.S., R.S.I., and M.Y. performed the field experiment. Y.S. analyzed the data. Y.S., R.S.I., K.K.S., and A.J.N. designed the project and wrote the manuscript with input from all co-authors.

## Ethics approval and consent to participate

Not applicable.

## Consent for publication

Not applicable.

## Competing interests

The authors declare that they have no competing interests.

## Supporting information

**Figure S1.**
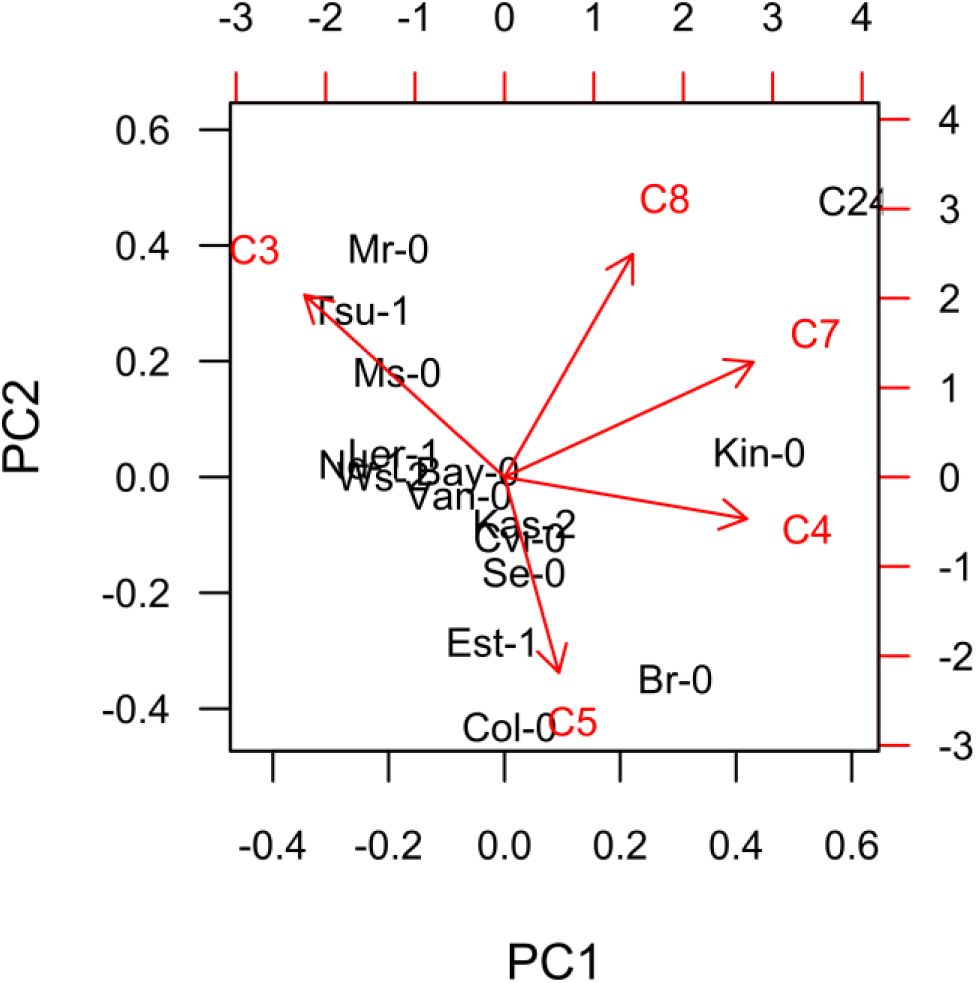
The first and second principal component (PC1 and PC2) summarizing the total amount (nmol/mg flesh weight) of C3-, C4-, C5-, C7-, and C8-Aliphatic glucosinolates for 17 accessions of *A. thaliana* (compiled from Chan et al. [34]). Arrows indicate contributions of each glucosinolate to PC1 and PC2.

**Figure S2.**
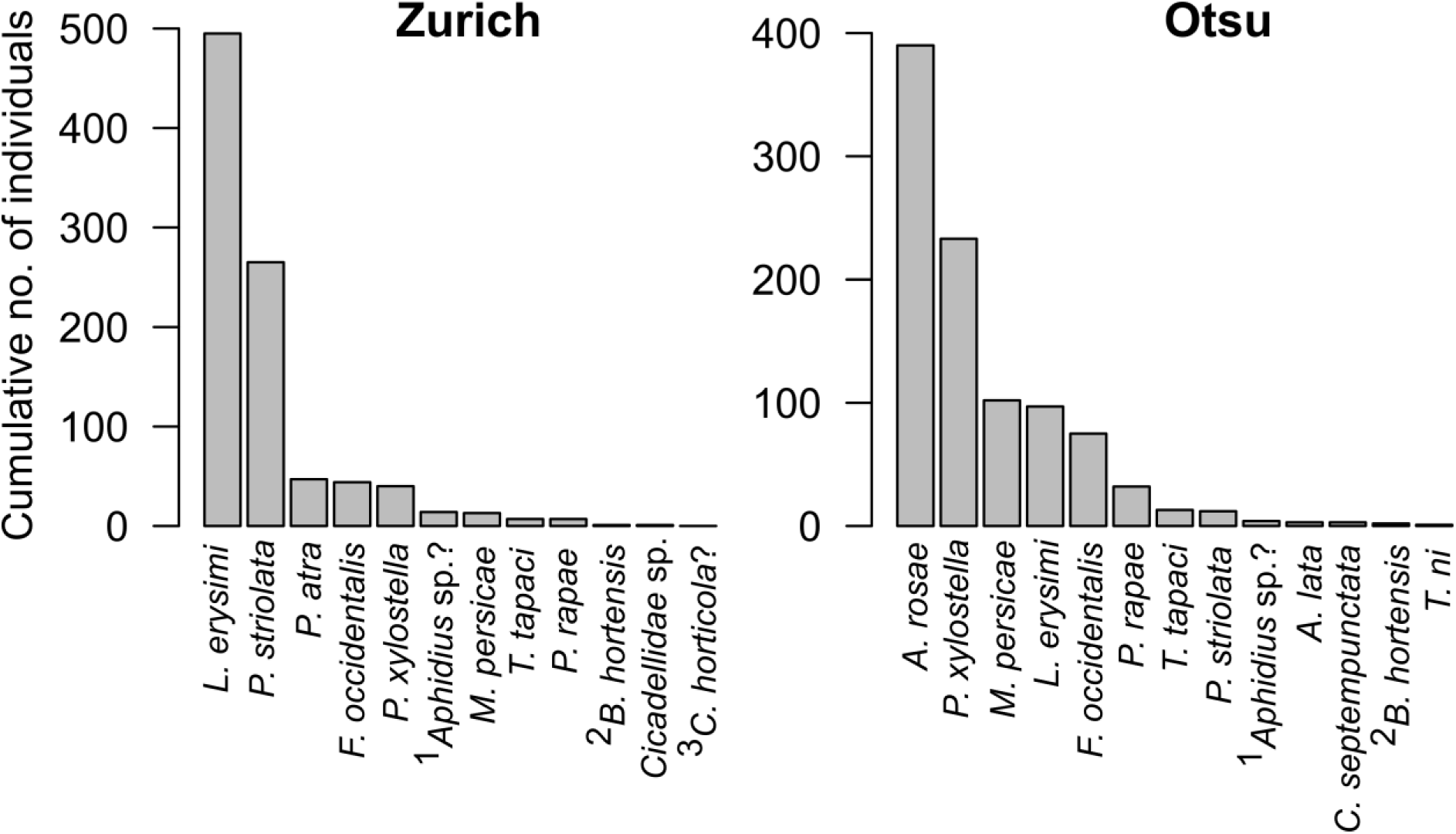
Cumulative number of each insect species in Zurich, Switzerland (left chart) and Otsu, Japan (right chart) throughout the experiments. See Table 2 for the name of the arthropod species. Notes: ^1^Total number of parasitoid wasps and mummified aphids; ^2^This species is a non-insect arthropod; ^3^Only a dwelling trace was observed.

**Table S1.**
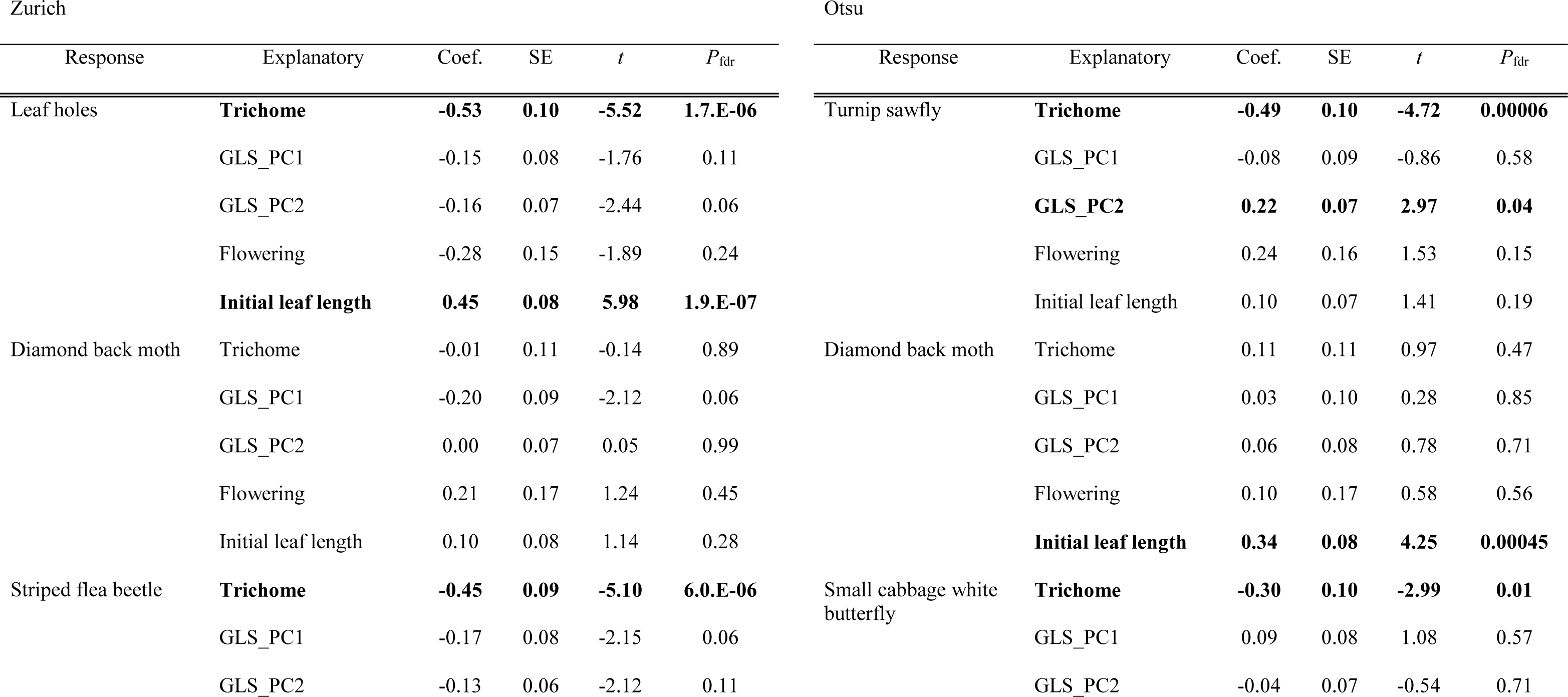
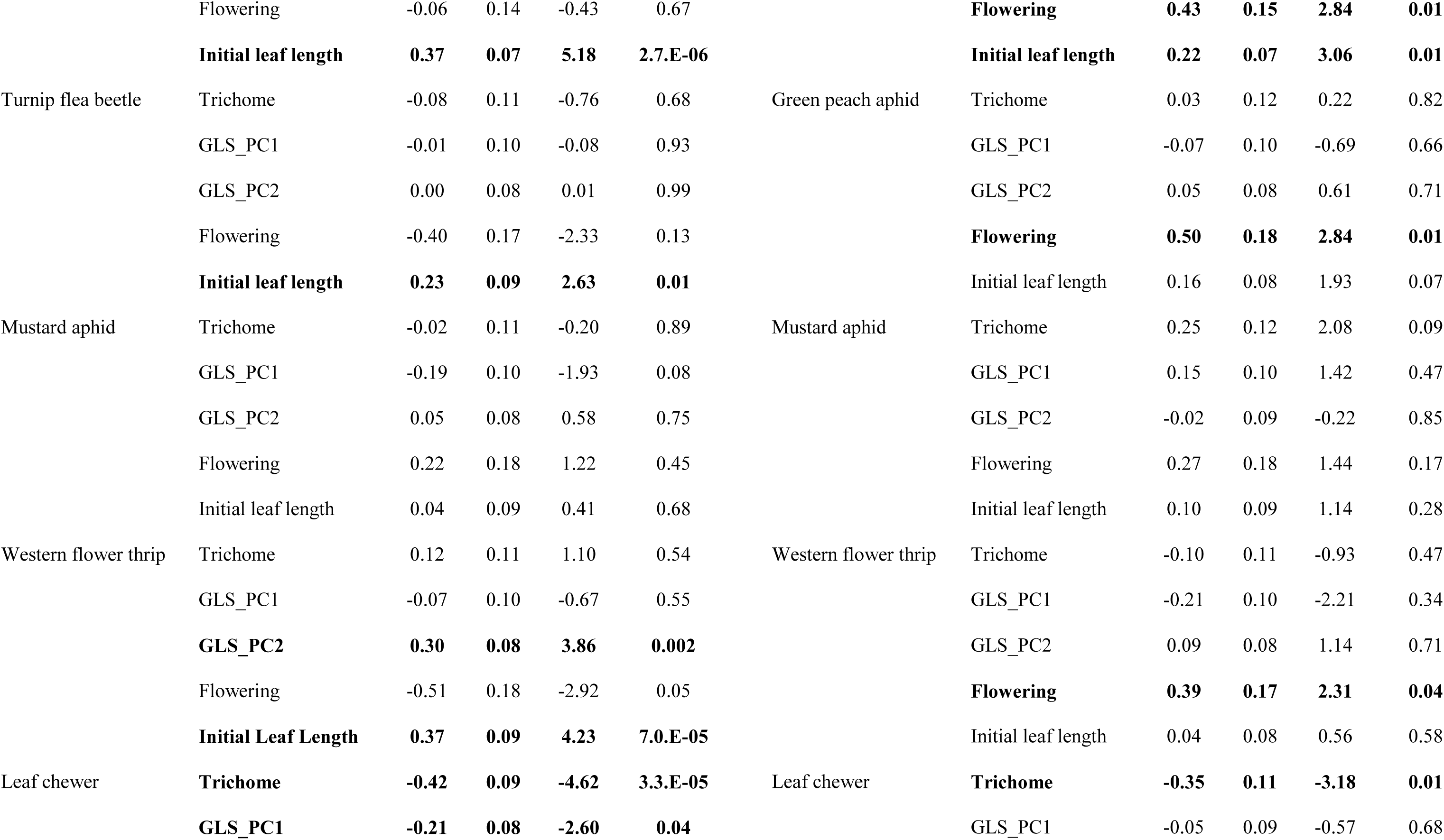
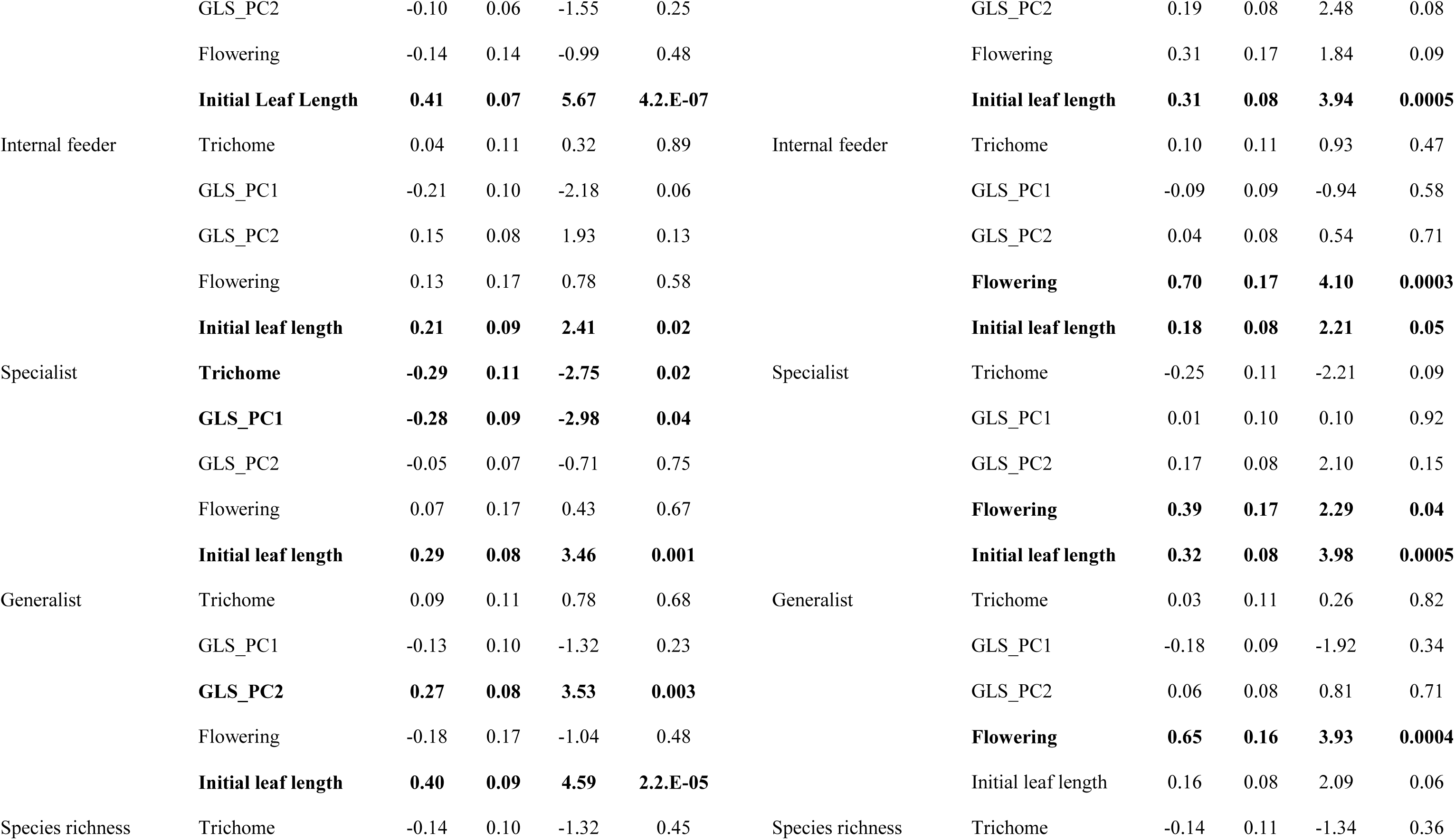
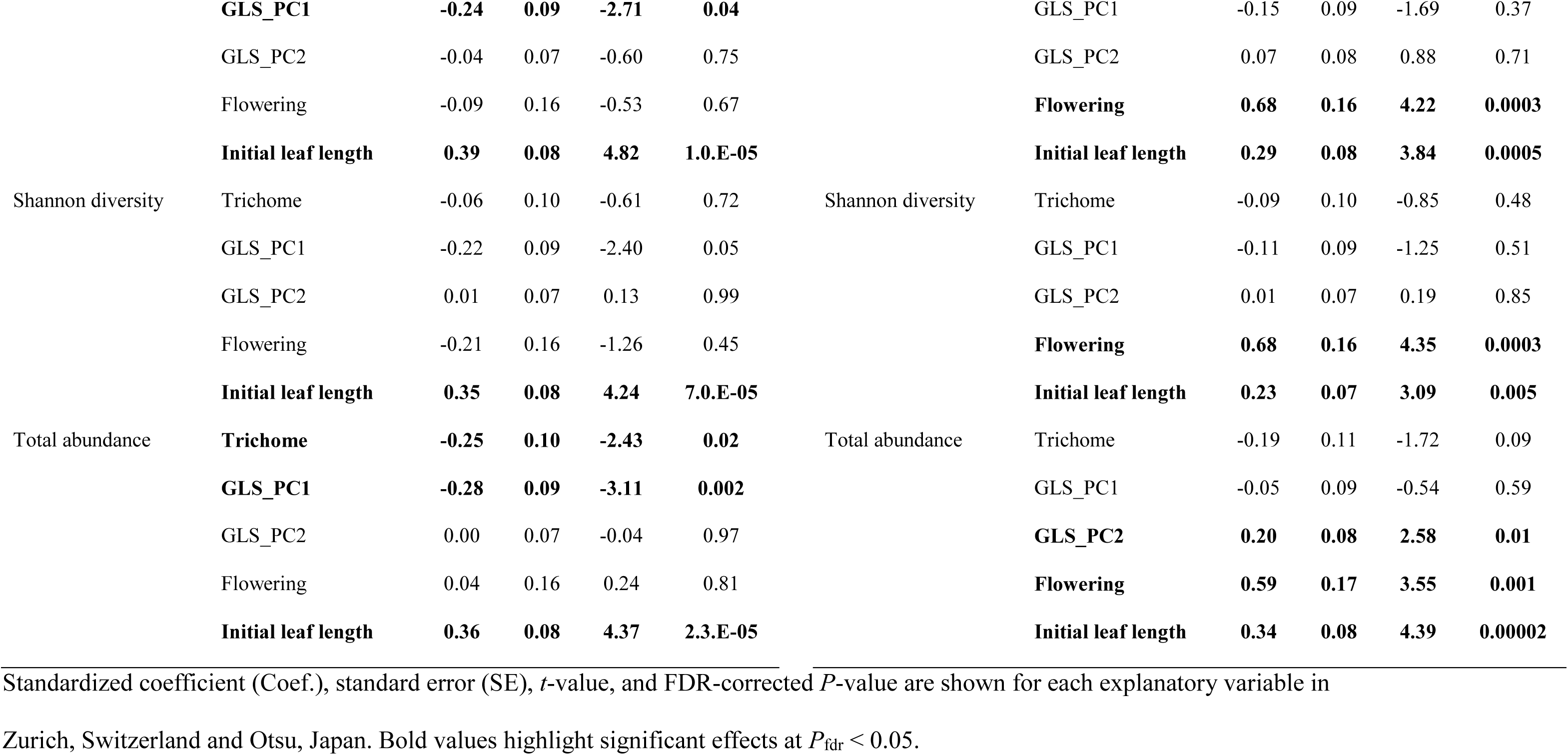
Effects of trichome density, the first and second principal component (PC1 and PC2) of aliphatic glucosinolates (GLSs), presence of flowering stem, and initial plant size on insect abundance and community composition among 17 natural accessions of *Arabidopsis thaliana*.

**Table S2.** Effects of trichome density, presence of flowering stem, and initial plant size on insect abundance and community composition in a comparison between glabrous mutants and their parental accessions.

| Zurich |  |  |  |  |  | Otsu |  |  |  |  |  |
| --- | --- | --- | --- | --- | --- | --- | --- | --- | --- | --- | --- |
| Response | Explanatory | Coef. | SE | *t* | *P*<sub>fd</sub> | Response | Explanatory | Coef. | SE | *t* | *P*<sub>fd</sub> |
| Leaf holes | **Trichome** | **-0.42** | **0.14** | **-3.05** | **0.027** | Turnip sawfly | **Trichome** | **-0.57** | **0.15** | **-3.94** | **0.002** |
|  | **Flowering** | **-1.18** | **0.27** | **-4.40** | **0.001** |  | Flowering | -0.42 | 0.29 | -1.46 | 0.397 |
|  | **Initial leaf length** | **0.42** | **0.13** | **3.14** | **0.021** |  | Initial leaf length | 0.23 | 0.16 | 1.44 | 0.318 |
| Leaf chewer | Trichome | -0.06 | 0.17 | -0.33 | 0.959 | Leaf chewer | **Trichome** | **-0.47** | **0.15** | **-3.07** | **0.009** |
|  | Flowering | -0.19 | 0.33 | -0.57 | 0.812 |  | Flowering | -0.43 | 0.30 | -1.40 | 0.397 |
|  | Initial leaf length | 0.39 | 0.16 | 2.38 | 0.062 |  | Initial leaf length | 0.26 | 0.17 | 1.55 | 0.318 |
| Specialist | Trichome | 0.01 | 0.17 | 0.05 | 0.959 | Specialist | **Trichome** | **-0.49** | **0.15** | **-3.16** | **0.009** |
|  | Flowering | 0.23 | 0.33 | 0.69 | 0.812 |  | Flowering | -0.40 | 0.30 | -1.31 | 0.397 |
|  | Initial leaf length | 0.32 | 0.16 | 1.95 | 0.068 |  | Initial leaf length | 0.26 | 0.17 | 1.58 | 0.318 |
| Species richness | Trichome | 0.01 | 0.17 | 0.05 | 0.959 | Species richness | Trichome | -0.24 | 0.17 | -1.37 | 0.214 |
|  | Flowering | 0.08 | 0.34 | 0.24 | 0.812 |  | Flowering | 0.12 | 0.34 | 0.36 | 0.722 |
|  | Initial leaf length | 0.32 | 0.17 | 1.89 | 0.068 |  | Initial leaf length | 0.08 | 0.19 | 0.40 | 0.689 |
| Shannon diversity | Trichome | -0.16 | 0.16 | -0.95 | 0.959 | Shannon diversity | Trichome | -0.17 | 0.18 | -0.97 | 0.339 |
|  | Flowering | -0.13 | 0.32 | -0.41 | 0.812 |  | Flowering | 0.14 | 0.35 | 0.40 | 0.722 |
|  | Initial leaf length | 0.36 | 0.16 | 2.25 | 0.062 |  | Initial leaf length | 0.13 | 0.19 | 0.71 | 0.582 |
| Total abundance | Trichome | 0.04 | 0.17 | 0.22 | 0.959 | Total abundance | **Trichome** | **-0.46** | **0.16** | **-2.86** | **0.011** |
|  | Flowering | 0.30 | 0.33 | 0.91 | 0.812 |  | Flowering | -0.16 | 0.32 | -0.52 | 0.722 |
|  | Initial leaf length | 0.33 | 0.16 | 2.00 | 0.068 |  | Initial leaf length | 0.18 | 0.17 | 1.03 | 0.468 |

Standardized coefficient (Coef.), standard error (SE), *t*-value, and FDR-corrected *P*-value are shown for each explanatory variable. Bold
values highlight significant effects at $P_{\text{fdr}} < 0.05$ . NA means no data available. The trichome density presents differences between the null
and hypomorphic mutants. Because the trichome density had a significant effect on flea beetles and sawflies (Table S1), we focused on
species, guild, and community indices comprising these two leaf chewers.

